# Antigen-level resolution of commensal-specific B cell responses enabled by phage-display screening and B cell tetramers

**DOI:** 10.1101/2023.10.13.562275

**Authors:** Sheenam Verma, Matthew J. Dufort, Tayla M. Olsen, Jasmine C. Labuda, Samantha Kimmel, Sam Scharffenberger, Andrew T. McGuire, Oliver J. Harrison

## Abstract

Induction of adaptive immune responses to commensal microbes is critical for tissue homeostasis, and perturbation of these responses is associated with multiple chronic inflammatory disorders. However, the mechanisms underlying the induction and regulation of mucosal B cells targeting commensal microbes remain poorly understood, in part due to a lack of tools to identify commensal-specific B cells *ex vivo*. To address this, we identified immunogenic protein epitopes recognized by Segmented Filamentous Bacteria (SFB)-specific serum antibodies using a whole-genome phage display screen and identified immunogenic proteins engaging IgA, IgG_1_ and IgG_2b_ responses. Using these antigens, we generated B cell tetramers to identify and track SFB-specific B cell responses in the gut associated lymphoid tissue during natural and *de novo* colonization. We revealed a compartmentalized response in SFB-specific B cell activation between Peyer’s patches and mesenteric lymph nodes, with a gradient of IgA, IgG_1_ and IgG_2b_ isotypes along the small intestine, and selective production of IgG_2b_ with the mesenteric lymph node chain. VDJ sequencing analyses and generation of SFB-specific monoclonal antibodies identified that somatic hypermutation drives affinity maturation to SFB derived antigens under homeostatic conditions. By combining phage display screening and B cell tetramer technologies, we now enable antigen-level based studies of immunity to intestinal microbes, which will advance our understanding of the ontogeny and function of commensal-specific B cell responses in tissue immunity, inflammation and repair.

## Introduction

Immunoglobulin A (IgA) plays a key role in immunity to pathogens, shapes early life interactions with the microbiota, and maintains microbiota diversity and compartmentalization throughout life^1–4^. In the gut, both T cell-independent (TI) and T cell-dependent (TD) mechanisms promote IgA production. TI IgA is considered to primarily recognize conserved microbial structures with relatively low affinity^5^. By contrast, TD IgA is strongly induced by mucus-resident and epithelial-adherent microbes and potently regulates mutualistic relationships with the microbiota^1,2^. TD-IgA induction occurs primarily in the Peyer’s patches of the small intestine, where activated B cells receive T cell help to undergo class-switch recombination (CSR), enter germinal centers (GC) and undergo somatic hypermutation (SHM) of immunoglobulin genes^6^. Commensal-specific IgG antibodies are also generated and can contribute to both host protection during systemic infection^7^, and immune regulation at mucosal sites by dampening responses to commensals^8^. How IgG responses to commensal microbes are mounted is poorly understood, in part because it is only recently appreciated that these responses can contribute to synergistic host-microbe interactions during homeostasis^8–10^. While homeostatic IgG production within the intestine is hypothesized to limit bacterial translocation and associated bacteremia, most of our understanding of commensal-specific IgG implicate these responses in disease pathogenesis as elevated levels of commensal-specific IgG antibodies and colonic IgG^+^ plasma cells are routinely observed in patients with severe Inflammatory Bowel Disease (IBD)^11,12^. A better understanding of how commensal-specific IgA and IgG responses are mounted and regulated is therefore important for studies of intestinal homeostasis, immunity, and inflammation.

Mucus-resident and epithelial-adherent microorganisms play a dominant rule in directing mucosal immune responses during homeostasis. In humans, non-human primates and rodents, epithelium-associated microorganisms influence the local and systemic activation of the immune system, and are implicated in both protection from pathogen infection, as well as exacerbation of tissue inflammation and autoimmunity. These species, including *Clostridiales spp*., *Candida spp*., *Akkermansia muciniphila* and *Helicobacter spp*., all interact closely with the intestinal epithelium^13–18^, a process thought to account for their ability to heighten tissue immunity^19^. Thus, investigating how commensal microbes that intimately engage with the intestinal epithelium influence host immunity is key to understanding not only tissue homeostasis, but also inflammation. In murine systems, a key example of these species is Segmented Filamentous Bacteria (SFB), which represents a prototypic commensal colonizing the gastrointestinal tract. Indeed, SFB colonization promotes accumulation of Th17 and Th1 cells in the small intestinal lamina propria, Th17 and T_FH_ cells in the Peyer’s patches, and production of T cell-dependent IgA^20–25^. SFB-specific Th17 cell responses have been associated with protection from gastrointestinal infection^20^, as well as disease exacerbation in the contexts of autoimmunity and autoinflammation^26–28^.

Current approaches for interrogating B cell responses to the microbiota rely on antibody-mediated immunoprecipitation of IgA/IgG-coated microbes, followed by 16S rRNA sequencing (IgA-seq/IgG-seq)^13,25,29,30^. Such approaches enable species-level resolution of immunogenicity but convey little information regarding the anatomic locale or cellular phenotype of B cells responding to individual microbes, nor the microbial antigens targeted. Conversely, elegant efforts to sequence B cell receptors (BCR) from gut B cells reveal a snapshot of B cell phenotype, but the immunogenic targets of these antibodies require further downstream identification using culture collections or antigen arrays^31–33^. This represents a formidable task given the complexity of the microbiota, with thousands of bacterial species each encoding hundreds to thousands of unique proteins, representing an overwhelming number of potential target antigens. We set out to develop a new high-throughput means to identify immunogenic antigens from commensal microbes, and subsequently conduct studies of commensal-specific IgA and IgG producing B cells at cellular resolution, which to date have not been feasible.

Phage display technology is a high-throughput screening technique used to identify ligands and binding partners for proteins and other macromolecules. This approach also enables identification of antibody binding to thousands of potential antigens in parallel, with previous successful employment identifying immunogenic antigens derived from pathogenic microbes^34,35^. Utilization of B cell tetramers is increasingly employed as an experimental approach to identify antigen-specific B cells in response to exposure to model antigens, vaccination and infectious pathogens. Indeed, use of immunogenic proteins from *Plasmodium spp.* and influenza A virus as B cell tetramers has been a key step in identifying aspects of host immunity to these pathogens within systemic and barrier tissues^36,37^. Broader application of these tools to immunogenic commensal microbes would enable breakthroughs in our understanding of the dynamic interactions between the immune system and the microbiota.

In this report, we now combine the cutting-edge approaches of phage-display screening and B cell tetramers, to identify immunogenic antigens and profile the ontogeny of B cells specific to commensal antigens *in vivo*. We generated a phage-display library derived from genomic DNA of Segmented Filamentous Bacteria to screen for immunogenic B cell antigens and discovered multiple SFB-derived antigens bound by distinct antibody isotypes. We subsequently generated B cell tetramers to directly identify SFB-specific B cells in the gut-associated lymphoid tissue (GALT) during vertical transmission and *de novo* colonization with SFB, revealing distinct B cell activation states in different anatomical sites. By performing BCR repertoire sequencing and clonal analyses, in conjunction with generating clonal lineages of SFB-specific antibodies, we directly demonstrate a trajectory of affinity maturation to a selected commensal antigen. This approach now enables the phenotypic interrogation of commensal-specific B cells across tissues and time, facilitating future studies of mucosal B cell biology in health and disease.

## Results

### Phage-display library screening identifies immunogenic commensal antigens

We sought to develop a system with which to track antigen-specific B cell responses to commensal microbes within the GALT. To do so, we selected Segmented Filamentous Bacteria as an immunostimulatory epithelium-adherent microbe capable of modulating host immunity, including induction of TD IgA production^21^. To identify immunogenic protein antigens recognized by SFB-specific B cells, we generated an SFB phage display library and antibody-based biopanning screen, an unbiased approach previously used to identify immunogenic B cell antigens in pathogenic microbes^34,35^. To this end, we cloned 200-600bp fragments of SFB genomic DNA into a phagemid library vector that fuses potential antigens to the M13 phage outer surface protein pIII^38^ **(Fig.1 A)**. The phagemid library was packaged to form infectious phage using pIII-deficient Hyperphage (M13K07ΔgIII), selecting for in-frame pIII-fusion ORFs by genetic complementation, and generating a phage library consisting of ~5×10^7^ putative SFB antigens **(Fig. 1A)** ^38,39^. Immobilized serum IgA and IgG antibodies were used to immunoprecipitate SFB epitope-bearing phage **(Fig. 1B)**. Specifically, serum from both germ-free and SFB-monocolonized mice was depleted of phage-reactive antibodies, and subsequently used to immunoprecipitate SFB antigen-bearing phage. Immunoprecipitated phage were released by trypsin digestion, and amplified by infection of F’ *E.coli*. To identify high affinity binders, we conducted 4 increasingly stringent biopanning cycles. After enrichment, we isolated 96 individual monoclonal phage, and using ELISA, retested individual phage for binding using serum from SFB monocolonized mice. This resulted in identification of 33 monoclonal phage (O.D. >0.5), with all identified phage encoding epitopes derived from the SFB genome **(Fig. 1C-E)**. Stochastic sampling of unbound phage identified both non-immunogenic inserts aligning to the SFB genome, but also inserts derived from the mouse genome, representing host gDNA content in fecal samples utilized for phagemid library preparation **(data not shown)**. Among the 33 monoclonal phage, we assigned antigens to 4 SFB-derived proteins **(Fig. 1E, S1A)**. Of the four identified antigens, SFB5460, SFB3310 and SFB15470 all had multiple repeat detections of a single immunogenic domain, whereas SFB3340 was identified to be targeted in 3 distinct N-terminal epitopes **(Fig. S1A)**. Additionally, while SFB5460 and SFB3340 were targeted by both IgA and IgG, SFB3310 and SFB15470 were bound selectively by IgG or IgA, respectively. **(Fig. 1E)**. As such, phage display screening represents a powerful experimental system with which to identify immunogenic commensal antigens, increasing the granularity of the targets of humoral immunity from the species level afforded by IgA/G-seq to that of individual immunogenic proteins. Strikingly, and validating our approach, among the immunogenic SFB antigens identified by phage display screening, SFB3340 has also been identified as a target of Th17 and T_FH_ cell responses elicited by SFB in the small intestinal lamina propria and Peyer’s patches, respectively^28^. Given the previous generation, and widespread employment of SFB3340-specific peptide:I-Ab tetramers and T cell receptor transgenic mice, we focused on this antigen to investigate commensal-specific B cells within systemic and gut associated lymphoid tissues.

**Figure 1:**
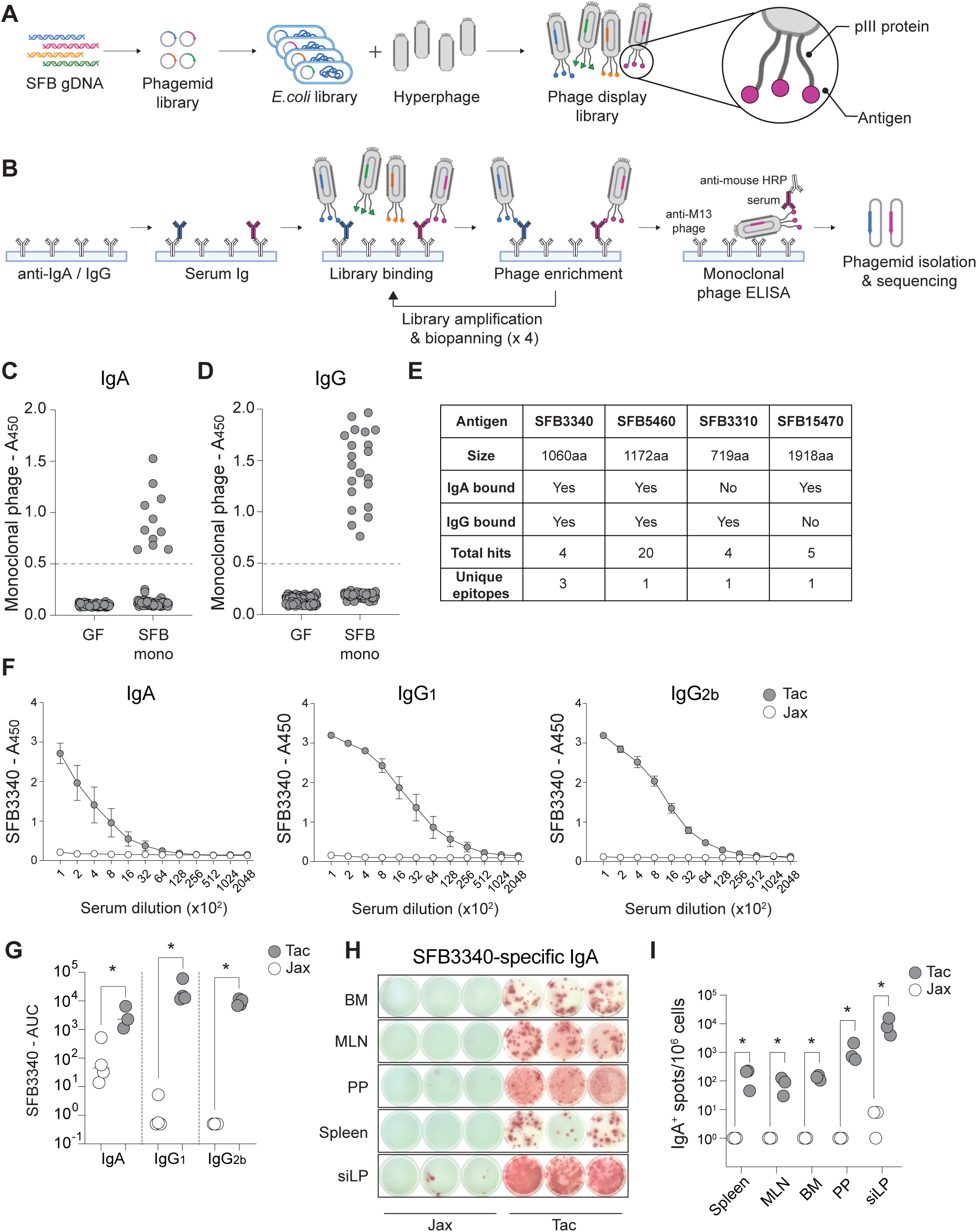
Phage display screening identifies immunogenic B cell antigens in Segmented Filamentous Bacteria. (A) Schematic of generation of SFB antigen phage display library. (B) Schematic of phage display biopanning using SFB antigen library and serum antibodies from SFB-monocolonized gnotobiotic mice. (C-D) Identification of immunogenic antigen hits using serum from germ-free or SFB monocolonized mice, assessed by binding of enriched monoclonal phage to immobilized (C) IgA, or (D), IgG. (E) Features of the 4 immunogenic SFB proteins identified by phage display. (F) SFB3340-specific ELISA of serum antibody reactivity by WT mice from Jackson Laboratories (Jax) or Taconic Farms (Tac). (G) SFB3340-specific ELISA binding represented as Area Under Curve (AUC). (H-I) Representative images and quantification of SFB3340-specific ELISpot IgA responses from leukocytes isolated from indicated tissues and stimulated with recombinant SFB3340. For phage display (A-E), data represent a single screen conducted with serum pooled from germ-free (n=3) or SFB-monocolonized (n=5) mice. For ELISA and ELISpot data (F-I), data represent at least 2 experiments with three to five mice per group. * p<0.05.

### Phage-display derived antigens identify commensal-specific B cells *ex vivo*

To validate SFB3340 as a target of mucosal B cells and investigate the extent of targeting of SFB3340 under homeostatic conditions in specified pathogen free (SPF) mice, we performed antigen-specific serum ELISA and ELISpot assays using tissues from naïve wildtype (WT) C57BL/6 (B6) mice naturally colonized (Taconic Farms - Tac) or not (Jackson Laboratories - Jax), with SFB. SFB3340-specific serum IgA was readily detectable in SFB-colonized Tac, but not Jax, B6 mice **(Fig. 1F**), confirming that SFB3340 is indeed targeted in specified pathogen free (SPF) mice containing a diverse commensal microbiota, and reactivity was not an artifact of gnotobiotic environs. Additionally, SFB3340-specific serum IgG_1_, and IgG_2b,_ but not IgM, IgG_2c_, IgG_3_ or IgE, were detected in SFB-colonized Tac B6 mice **(Fig. 1F,G**, **S1B)**. In line with reports that SFB-specific IgA responses are entirely T-dependent^21,25^, SFB3340-specific IgA, IgG_1_ and IgG_2b_ were undetectable in *Tcrbd*^−/−^ mice **(Fig. S1C)**. Furthermore, these responses were undetectable in both *Icos^−/−^* and *Bcl6*^1′CD^^4^ mice, demonstrating a requirement for T_FH_ cells^40,41^ **(Fig. S1D-E)**. To identify the distribution of SFB3340-specific B cells across tissues, we used antigen-specific ELISpot assays. SFB3340-specific B cell responses producing IgA, IgG_1_ or IgG_2b_ were identified in the spleen, Peyer’s patches (PP), mesenteric lymph nodes (MLN), small intestinal lamina propria (siLP) and bone marrow (BM) of Taconic, but not Jax mice **(Fig. 1H-I, S1F-G)**. Together, SFB3340 is the target of T_FH_-dependent IgA, IgG_1_ and IgG_2b_ producing B cells and serves as a tool to investigate the induction of commensal-specific B cells responding to an endogenous antigen under homeostatic conditions.

### B cell tetramers enable identification of commensal-specific B cells *ex vivo*

Identification of immunogenic commensal antigens provides a unique opportunity to address key unanswered questions regarding mucosal immunity that cannot be addressed without direct identification of commensal-specific B cells. B cell antigen tetramers are powerful tools that enable analysis of cellular phenotype, isotype, kinetics and polyclonal repertoire of antigen-specific B cells. We extended these utilities to study SFB-specific B cells in the GALT, by adopting techniques used to study protein immunization and pathogen infection^36,42^. To this end, we generated phycoerythrin (PE)-conjugated B cell tetramers containing the majority of the SFB3340 protein, lacking the putative N-terminal transmembrane domain **(Fig. S2A)**. In all experiments, isolated leukocytes were initially stained with a decoy reagent, to identify and exclude cells binding structural components of the tetramer^42^ **(Fig. S2A)**. Cells were subsequently stained with SFB3340 tetramers and fluorescent antibodies for analysis by multiparameter flow cytometry, and after excluding non-B cells and doublets, SFB3340^+^Decoy^−^ B cells were identified among B220^+^ and B220^low^CD138^+^ cells **(Fig. S2B)**. In naïve Jax B6 mice that are not colonized by SFB, the few SFB3340^+^ B cells identified within Peyer’s patches were naïve follicular B cells (IgD^+^CD38^+^GL-7^−^FAS^−^), representing the naïve immune repertoire **(Fig. 2A-C)**. By contrast, in the Peyer’s patches of SFB-colonized Tac B6 mice, we identified SFB3340^+^ B cells with an activated phenotype. Specifically, we identified 77±6% of SFB3340^+^ B cells to be IgM^−^IgD^−^ class-switched (swIg^+^), compared to 14±4% of polyclonal B cells in Tac B6 mice **(Fig. 2C)**. swIg^+^ SFB3340^+^ B cells were not detected in Peyer’s patches of Jax B6 mice, consistent with a requirement of antigen for B cell activation **(Fig. 2C)**. Furthermore, nearly all swIg^+^ SFB3340^+^ B cells were FAS^+^CD38^−^ germinal center (GC) B cells, with few if any FAS^−^CD38^+^ memory B cell (MBC) detectable in steady state mice **(Fig. 2D-E)**. By contrast, MBC were detectable among the polyclonal repertoire of swIg^+^ Peyer’s patch B cells **(Fig. 2D-E)**. A similarly polarized Peyer’s patch GC B cell response to SFB3340 was also detected in SFB monocolonized, but not germ-free mice, and in Jax B6 mice *de novo* colonized with feces from SFB-monocolonized mice **(Fig. 2F-G)**. Thus, our data demonstrate that SFB3340 B cell tetramers now enable direct identification of SFB-specific B cell activation, and that SFB colonization is necessary and sufficient to elicit induction of SFB3340^+^ GC B cell responses within Peyer’s patches.

**Figure 2:**
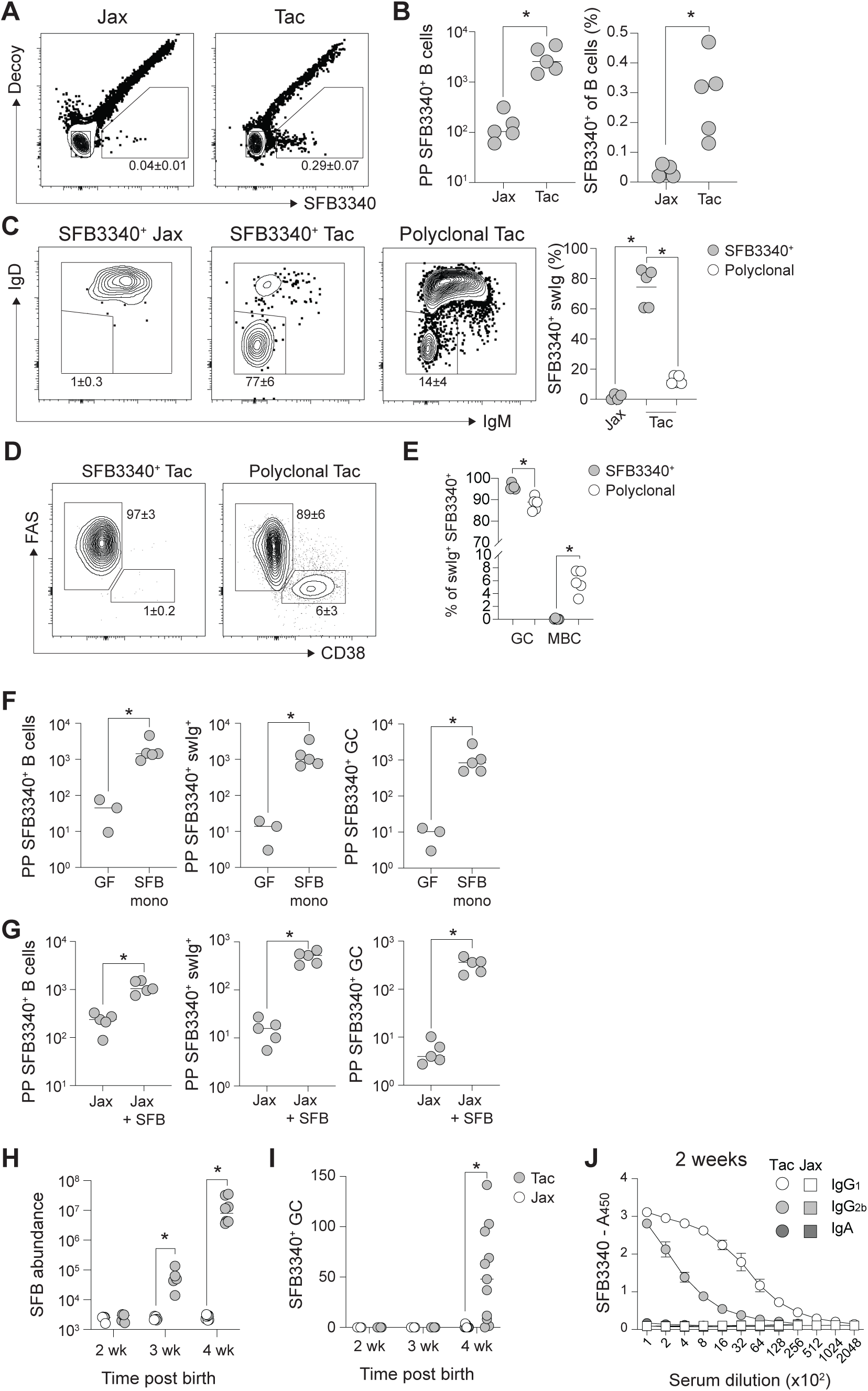
Natural SFB colonization promotes SFB3340-specific GC B cell responses in Peyer’s patches. (A) Representative contour plots of SFB3340 B cell tetramer and decoy reagent staining in Peyer’s patch B cells. (B) Total number and relative frequency of SFB3340^+^ B cells in Peyer’s patches of WT mice from Jackson Laboratories (Jax) or Taconic Farms (Tac). (C) Representative contour plots of IgM and IgD expression and frequencies of swIg^+^ SFB3340^+^ B cells in Peyer’s patches in WT B6 mice, or polyclonal SFB3340^−^ B cells in Taconic B6 WT mice. (D-E) Representative contour plots and relative frequencies of GC and MBC populations on SFB3340^+^ or polyclonal B cells in Peyer’s patches of SFB-colonized Taconic B6 WT mice. (F) Total numbers of SFB3340^+^ B cells, swIg^+^ B cells, and GC B cells in germ-free and SFB-monocolonized mice. (G) Total numbers of SFB3340^+^ B cells, swIg^+^ B cells and GC B cells in Jax B6 mice colonized, or not, with fecal contents from SFB-monocolonized gnotobiotic mice. (H) Quantification of SFB load in fecal contents assessed by qPCR, (I), frequencies of SFB3340^+^ GC in Peyer’s patches, and (J), serum antibody titers of SFB3340-specific antibodies in pups of Jax and Tac B6 breeding pairs at indicated timepoints post birth. Data represent at least 2 experiments with three to five mice per group. * p<0.05.

Recent studies revealed that a substantial proportion of homeostatic intestinal IgA^+^ cells in adult mice arise early in life^43^. To investigate the dynamics of early life SFB3340-specific B cell activation during natural colonization by vertical transmission, we conducted parallel timed breeding of Jax B6 and Tac B6 mice. As previously described^44^, SFB colonization was undetectable by qPCR in either strain at 2 weeks of age, but readily detectable in Tac B6 mice by 3 weeks of age, and reached levels observed in adult mice by 4 weeks post-birth **(Fig. 2H)**. Induction of SFB3340^+^ B cell activation in Peyer’s Patches was detectable by 4 weeks of age **(Fig. 2I)**. Strikingly however, and consistent with reports of transplacental transfer of IgG_1_ and IgG_2b_ ^8^, serum antibodies specific to SFB3340 were already detectable in serum by 2 weeks of age, significantly preceding both SFB colonization and SFB3340^+^ B cell activation **(Fig. 2J)**. SFB3340- specific IgA was not detected at this timepoint, in line with the observation that IgA is not trans-placentally transferred, nor integrated into the circulation following ingestion. As such, SFB3340-specific B cell tetramers enable discrimination of host B cells and maternally derived antibody responses during early life.

### Compartmentalized induction of distinct isotypes by SFB3340^+^ B cells

Recent studies identified compartmentalized drainage of the intestine by distinct lymph nodes within the gut draining lymph node chain^45–47^. Functionally, this process results in specialization of individual gut LNs in priming tolerogenic or pro-inflammatory T cell responses and the subsequent accumulation of regulatory or effector T cell populations in distinct gut segments^45–47^. We investigated whether a similar gradient of specialized differentiation operates for commensal-specific B cells within Peyer’s patches. Strikingly, analysis of Peyer’s patches from duodenal, jejunal and ileal regions of Tac mice demonstrated clear compartmentalization of antibody isotype production by SFB3340^+^ B cells. While GC B cell frequency and low IgG_1_ expression among swIg^+^ SFB3340^+^ B cells was stable across Peyer’s patches from distinct intestinal regions **(Fig. 3A-B)**, proximal Peyer’s patches (duodenal and jejunal) were highly enriched in IgG_2b_^+^ GC B cells, with a significantly lower frequency in ileal Peyer’s patches **(Fig. 3B)**. By contrast, IgA^+^ GC B cell frequencies were low in proximal Peyer’s patches, but represented the dominant antibodies produced in ileal Peyer’s patches **(Fig. 3B)**. To further determine how anatomic locale governs SFB-specific B cell differentiation, we compared total Peyer’s patches with cells isolated from MLN of adult Taconic B6 mice. While swIg^+^ SFB3340^+^ B cell responses were detectable in both tissues, the frequency of swIg^+^ SFB3340^+^ B cells was significantly lower in MLN **(Fig. 3C)**. Strikingly, though IgA^+^ GC B cells were readily detected in the polyclonal B cell population, MLN-derived SFB3340^+^ GC B cells did not produce IgA, and instead almost exclusively secreted IgG_2b_ antibodies **(Fig. 3D-F)**. Therefore, using B cell tetramers, we now demonstrate a clear anatomical compartmentalization of SFB-specific B cell effector functions across the Peyer’s patches and within the MLN.

**Figure 3:**
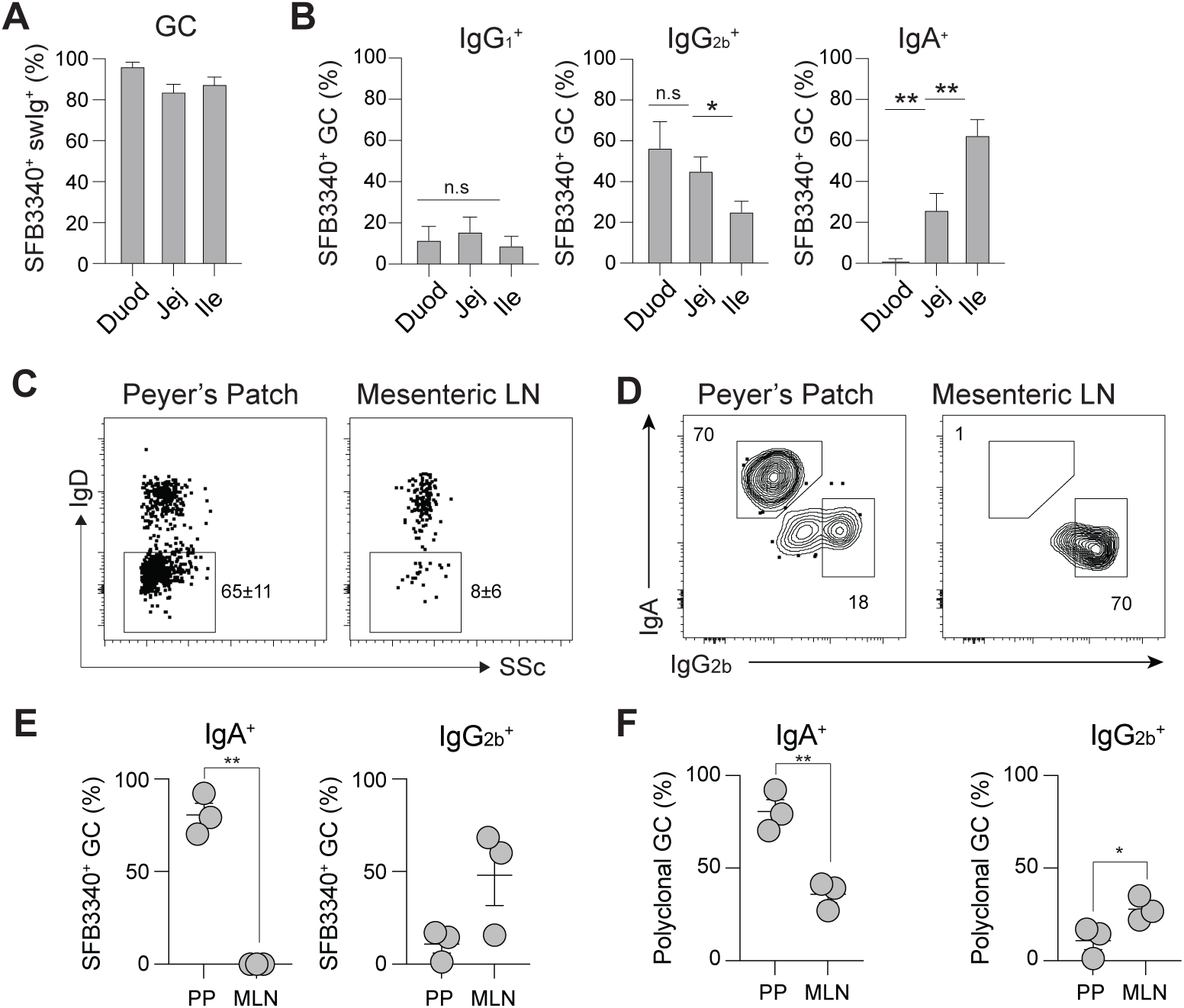
Compartmentalized induction of distinct isotypes to SFB3340 antigen. Peyer’s patches were isolated from distinct intestinal regions of Tac B6 mice and assayed for SFB3340^+^ B cell phenotype. (A) Relative frequencies of GC B cells among swIg^+^ SFB3340^+^ B cells, and (B) relative frequencies of isotype expressing SFB3340^+^ GC B cells in distinct intestinal sections. (C) Representative flow cytometry plots and frequencies of swIg^+^ SFB3340^+^ B cells in total gut Peyer’s patches or mesenteric lymph nodes. (D) Representative flow cytometry plots and (E) frequencies of IgA^+^ and IgG_2b_^+^ SFB3340^+^ GC B cells in Peyer’s patches and MLN. (F) Relative frequencies of IgA^+^ and IgG_2b+_ polyclonal GC B cells in Peyer’s patches and MLN. Data represents one of two independent experiments with three to five mice per group. * p<0.05, ** <0.01.

### Kinetic analyses of SFB-specific B cell responses during *de novo* colonization

To gain a deeper understanding of the kinetics and phenotype of SFB-specific B cells induced in GALT without the influence of maternal antibodies acquired during vertical transmission^8^, we introduced SFB into adult naïve Jax B6 mice by oral gavage. We used filtered fecal contents from SFB-colonized *Rag2*^−/−^*Il2rg*^−/−^ mice, which harbor ~10 fold enriched fecal SFB compared to WT animals, mimicking a previously described strategy using enriched-SFB (eSFB) flora from lymphocyte-deficient *Rag2*^−/−^*Il23r*^−/−^ mice^48^. This approach also benefits from a lack of donor-derived immunoglobulins that coat the fecal contents of WT mice, that could influence *de novo* colonization by SFB. We tracked the kinetics and phenotype of SFB3340-specific B cells in GALT in naïve mice and mice 7, 14, 21 and 28 days post-colonization (d.p.c.). In concordance with the rate of engraftment and establishment of stable SFB colonization **(Fig. S4A)**, activated SFB3340^+^ B cells were detectable by 14 d.p.c., with maximal engagement of swIg^+^ SFB3340^+^ B cells at 28 d.p.c. **(Fig. 4A-B)**. At timepoints prior to 14 d.p.c., numbers of SFB3340^+^ B cells were rare, precluding further investigation. By contrast, SFB3340-specific GC B cells were readily detectable from 14 d.p.c. onwards **(Fig. 4C-E)**. Kinetic analyses of SFB3340^+^ B cell isotype expression identified a stable distribution of IgA^+^, IgG_1_^+^ and IgG_2b_^+^ cells in the Peyer’s patches **(Fig. 4F-H)**. Over this time period, IgA and IgG_2b_^+^ B cells were maintained at stable relative frequencies, suggesting that IgA^+^ B cells were not outcompeted in gut GC responses, in line with recent work demonstrating that heightened IgA BCR-signaling promotes the survival of germinal center B cells in Peyer’s patches^49^.

**Figure 4:**
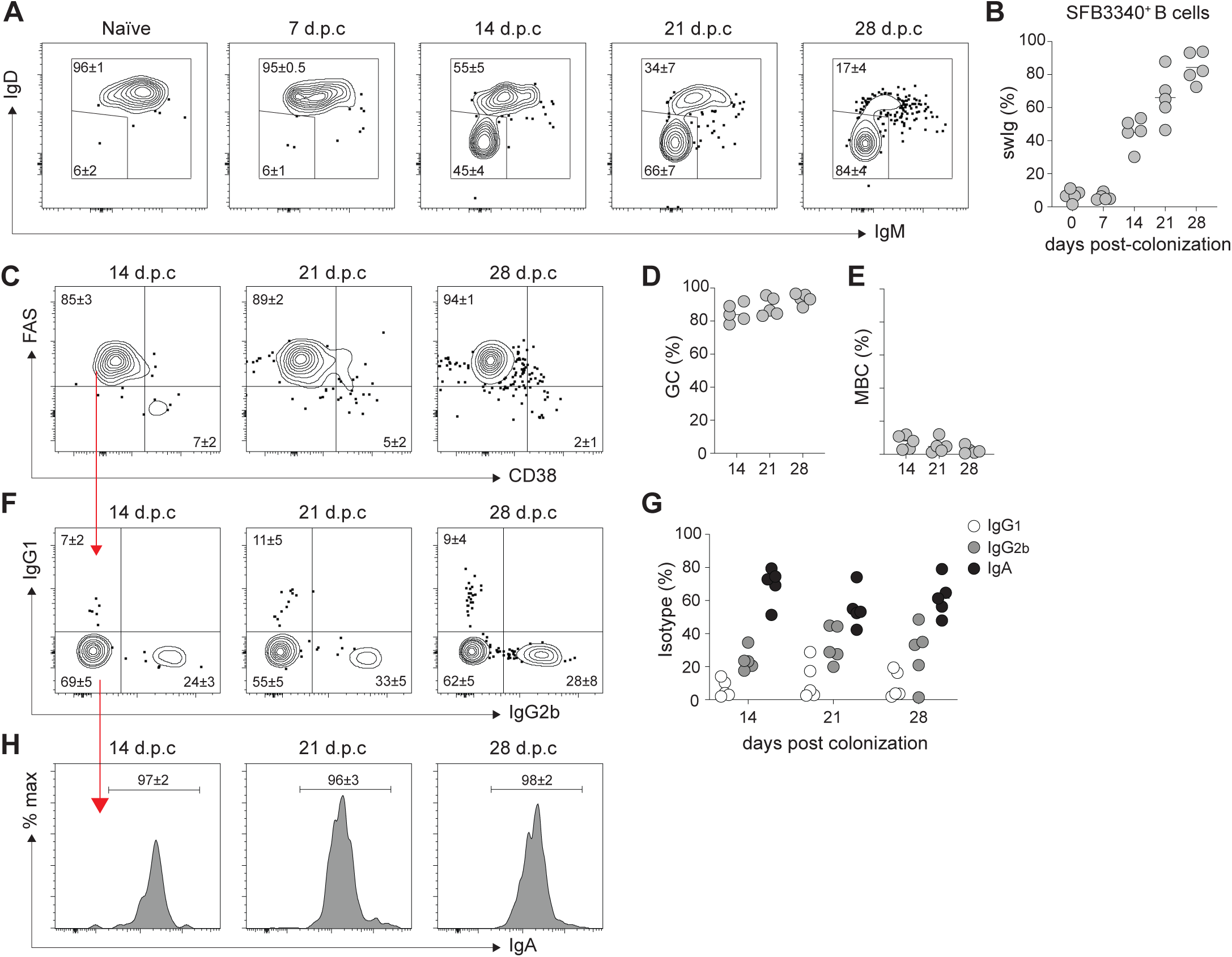
De novo colonization with SFB drives SFB3340-specific GC B cell induction with a broad range of antibody isotypes. (A-G) Jax B6 WT mice were colonized with SFB by oral gavage of fecal contents from SFB^+^ *Rag2*^−/−^*Il2rg*^−/−^ mice prior to analysis at 7, 14, 21 and 28 days post-colonization (d.p.c). (A) Representative contour plots of IgM and IgD expression by SFB3340^+^ Peyer’s patch B cells at indicated timepoints. (B) Relative frequencies of swIg^+^ SFB3340^+^ B cells in Peyer’s patches following SFB colonization. (C) Representative contour plots and (D-E) relative frequencies of GC (FAS^+^CD38^−^) and MBC (CD38^+^FAS^−^) SFB3340^+^ swIg^+^ B cells at 14, 21 and 28 d.p.c. (F & H) Representative contour plots and (G) relative frequencies of antibody isotype expression by SFB3340^+^ Peyer’s patch GC B cells following SFB colonization. Data represent at least 3 independent experiments with 4-5 mice per group.

### Affinity-driven commensal-specific B cell selection in Peyer’s patch germinal centers

Identification and isolation of single antigen-specific B cells from constitutively active germinal centers within Peyer’s patches enabled us to investigate the selection of gut GC B cells with unprecedented specificity and resolution. To date, the antigenic targets of Peyer’s patch B cells undergoing somatic hypermutation and affinity maturation remain to be determined *a priori*, requiring generation of monoclonal antibodies and subsequent identification of immunogenic microbes using microbial culture collections or co-opting pathogen-derived glycan arrays^31,32^. As such, we directly investigated the BCR repertoire usage and selection of gut B cells specific to SFB3340 using single cell RNA / VDJ-sequencing, and identified V(D)J segment usage, clonality and somatic hypermutation (SHM) profiles across SFB3340^+^ B cells following *de novo* colonization with SFB. To profile the repertoire of multiple mice, total PP cells were isolated from 10 individual WT mice at 21 d.p.c, stained using oligo-hash tagged and fluorescent antibodies, decoy and SFB3340 tetramer reagents and subsequently pooled. SFB3340^+^ B cells from the pooled sample were isolated by FACS and subjected to single cell RNA / VDJ sequencing.

Deconvolution of oligo-hashtagged antibodies demonstrated that individual mice were evenly represented amongst high quality cells, with phenotypic diversity mirroring our cytometric data **(Fig. S5A)**. We identified UMAP clusters based on marker gene expression, including naïve (*Ighd*), GC (*Aicda*) and a small population of antibody-secreting cells (ASC, (*Prdm1*)) **(Fig. 5A-C)**. Amongst these cellular populations, antibody isotype identified by scRNA-seq also correlated with cytometric analysis, with a majority of IgA and IgG_2b_-producing cells within the GC and ASC populations **(Fig. 4F-H**, **5B)**. To investigate BCR repertoire usage by SFB3340^+^ B cells, we mapped the V_H_ repertoire of productive V(D)J usage across the V_H_ locus and identified public clonotypes that recurred in GC B cells of multiple mice. We focused on the widespread utilization of two recurrent clonotypes (RC), V_H_1-76RC and V_H_1-53RC, with multiple mice utilizing a recurring clonotype (RC) of either, or often both, of these two segments. To determine the relationship between clonal selection and affinity maturation of SFB3340^+^ B cells using public V_H_1-76(RC) and V_H_1-53(RC) clonotypes, we reconstructed BCR evolution for these clonotypes from individual mice and generated phylogenetic trees of inferred SHM acquisition **(Fig. 5D-E)**. To determine whether acquisition of somatic hypermutations contributed to affinity maturation of SFB3340-specific responses, V_H_1-53RC-V_k_4-58J_k_1 and V_H_1-76RC-V_k_5-43J_k_2 antibody families and their unmutated ancestors (UA) were expressed as monoclonal antibodies in a human IgG_1_ backbone and binding to SFB3340 was assessed. As expected, all recombinant monoclonals from both V_H_1-53RC-V_k_4-58J_k_1 and V_H_1-76RC-V_k_5-43J_k_2 families bound SFB3340 protein **(Fig. 5F-I)**. ELISA and biolayer-interferometry (BLI)-based quantification of binding demonstrated a clear increase in binding by both V_H_1-53RC-V_k_4-58J_k_1 and V_H_1-76RC-V_k_5-43J_k_2 antibody lineages bearing somatic mutations **(Fig. 5F-I)**. Furthermore, reverting all IgH and IgL mutations to germline greatly reduced the binding of UA antibodies to SFB3340 **(Fig. 5F-I)**, strongly supporting that affinity maturation to SFB3340 occurs in PP GCs. Furthermore, affinity maturation was in part mediated by mutations accumulated within the IgH CDR3, as experimental introduction of acquired CDR3 mutations into UA IgH sequences increased binding even in the presence of UA IgL chains, demonstrating that CDR3 binding contributed to affinity maturation of SFB3340-specific antibody lineages **(Fig. 5J-K)**. Importantly, recombinant SFB3340-specific antibodies did not bind to irrelevant antigens that are common targets of polyreactive antibodies including Insulin, DNA, LPS, and ovalbumin ^5^ **(Fig. S5B)**. To demonstrate that recombinant antibodies identified by isolation of SFB3340^+^ B cells bind SFB3340 antigen in its native conformation, we conducted fluorescence microscopy on ileal tissue sections. Indeed, SFB3340-specific monoclonal antibody binding identified SFB in close physical proximity to the intestinal epithelium **(Fig. 5L),** confirming that these antibodies recognize SFB *in situ* within ileal tissues. Thus, B cell tetramers enable highly specific isolation of commensal-specific B cells, and SFB3340-specific B cell in Peyer’s patches undergo somatic hypermutation and affinity maturation during *de novo* colonization, as demonstrated by increasing binding affinity in antibodies bearing multiple mutations.

**Figure 5:**
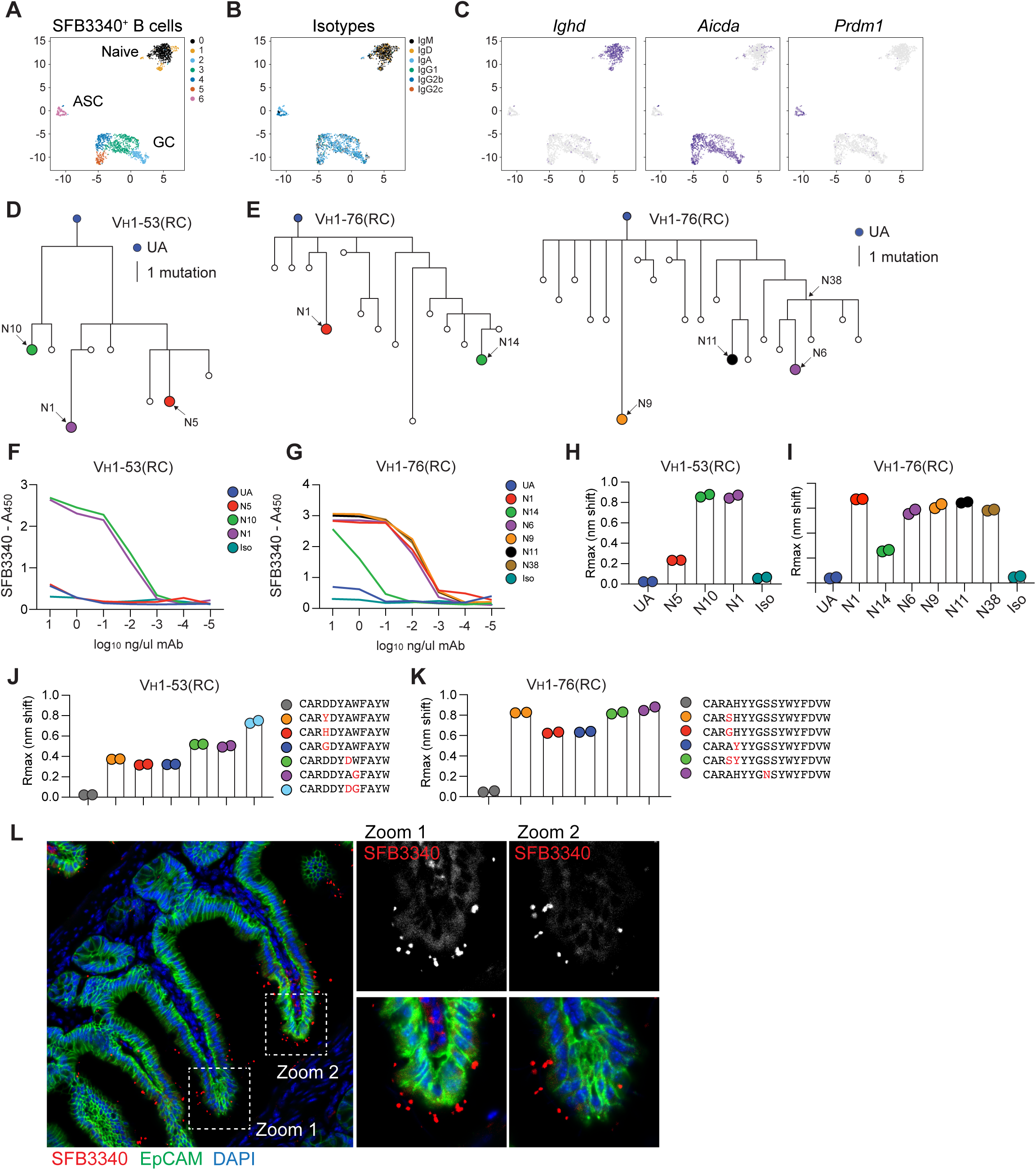
SFB3340^+^ GC B cells undergo somatic hypermutation and affinity maturation in Peyer’s patches under homeostatic conditions. (A-B) SFB3340^+^ B cells were isolated by FACS from the Peyer’s patches of WT B6 mice *de novo* colonized with SFB. Peyer’s patches from 10 mice were stained with SFB3340 tetramer and oligo-hashtagged antibodies, before pooling, and isolation of SFB3340^+^ B cells by FACS prior to scRNA-seq using the 10X platform. UMAP plots of (A) scRNA-seq clusters, (B) antibody isotype transcripts and (C) curated marker genes used to identify naïve, GC and antibody-secreting cell clusters. (D-E) Phylogenetic trees of reconstructed evolution of V_H_1-76(RC) and V_H_1-53(RC) antibody lineages identified from SFB3340^+^ B cells of individual mice. (F-G) ELISA based antibody binding of recombinant antibody lineages to recombinant SFB3340. BLI based antibody binding metrics of recombinant mAb from V_H_1-76(RC) and V_H_1-53(RC) antibody lineages and (J-K) UA antibodies bearing CDR3 mutations. (L) Confocal microscopy imaging of V_H_1-76(RC) N1 mAb staining of ileal tissue from an SFB-colonized *Rag2*^−/−^*Ilrg2*^−/−^ mouse.

## Discussion

Host immune responses elicited by commensal microbes influence tissue immunity, inflammation and repair, but also disease manifestation, severity and therapeutic efficacy^9,50^. Cognate T cell responses to bacterial components of the gut and skin microbiome play essential roles in maintenance and restoration of tissue integrity during infection and injury. While antibody responses to the microbiota are critical for microbiota containment, diversification, and gut function, our understanding of commensal-specific B cell responses is less developed due to our inability to track cognate responses beyond the community level of gut bacteria, much less the resolution of individual microbes. Although identification of immunogenic commensal microbes using immunoglobulin-based enrichment and subsequent 16S rRNA sequencing has shed light on the identity of immunogenic and disease-influencing microbes, identifying the ontogeny and cellular phenotype of B cells producing commensal-specific antibodies remains challenging. Our study details technical innovations that enable mechanistic investigation of antigen-specific B cells within barrier tissues, utilizing high-throughput screening to identify immunogenic commensal antigens, and converting these antigens into experimental reagents with which to identify cognate B cell responses to gut microbes.

As proof-of-concept, phage-display screening identified immunogenic antigens derived from Segmented Filamentous Bacteria, and significantly increased the resolution of antigenicity from an individual microbe to a handful of protein antigens. In itself this methodology enables use of antigen-specific ELISA or ELISpot to study immunity to a given microbe of interest and allows screening of humoral responses across experimental cohorts. While a powerful approach for antigen identification, phage display is restricted by the necessity for DNA encoded antigens, therefore limiting its utility for discovery of carbohydrate or lipid antigens. Similarly, due to the size restriction of gDNA inserts used during library generation, detection of complex antigens derived from multimeric or conformational epitopes also represents a technical limitation of the approach. Despite these considerations, phage display screening is a powerful technique to identify immunogenic antigens derived from commensal microbes. Indeed, although here demonstration of the utility of phage display screening focused on an intestinal bacterium, this approach can be readily adapted to study non-bacterial components of the microbiota, including fungi, viruses, helminths, protists as well as our genome-encoded symbionts, endogenous retroviruses. Furthermore, this approach could be employed in studies of immunity in other barrier tissues colonized by commensal microbes, including the skin, oral mucosa and female reproductive tract.

To further investigate the ontogeny, kinetics, phenotype and anatomic localization of SFB-specific B cells, we generated B cell tetramers from SFB3340, and established methods to identify SFB-specific cells *ex vivo* at mucosal sites, bridging the current divide between bacteria-level resolution of immunity enabled by IgA/G-seq, and the antibody-specificity resulting from sequencing of GALT-derived BCRs. Notably, this antigen-level resolution revealed previously undescribed compartmentalization of microbiota-specific antibody isotype production along the length of the small intestine, a dichotomy in isotype production between mesenteric lymph nodes and Peyer’s patches, and the kinetics of early life B cell responses to commensal microbes previously obfuscated by inheritance of maternal-derived antibodies. Specifically, we identified a clear anatomical compartmentalization of B cell effector functions with IgA^+^ SFB3340^+^ GC responses enriched in distal PP, and strong enrichment of IgG isotypes within proximal PP and MLN. These discoveries are in line with a model in which microbial components within the MLN might indicate bacterial translocation or infection, wherein the MLN would represent the last component of the mucosal firewall prior to systemic bacterial exposure. In this case, production of IgG isotypes would favor rapid opsonization and clearance of invading microbes. Investigating how the environmental milieu and local T cell populations within distinct PP and MLN influence selection of IgA^+^ and IgG^+^ B cells in gut germinal centers is of great interest. It is reasonable to hypothesize that this gradient in B cell effector function across intestinal regions occurs as a result of preferential colonization of the ileum by SFB, but also by local enrichment of factors promoting IgA induction, including Serum Amyloid A (SAA), implicated in licensing Th17 cell effector function along the length of the small intestinal lamina propria^48^, likely due to its role as a retinoic acid carrier^51^. In addition to detection of B cell activation across tissue sites, commensal-specific B cell tetramers enabled detection of early life B cell activation and the study of transgenerational immunity to commensal microbes in neonatal and adult mice. Looking forward, these tools will enable future investigation of early life humoral responses to natural colonization currently requiring models of maternal B cell/antibody deficiency and/or cross-fostering-based approaches^8^.

Finally, using *a priori* selection of SFB3340-specific B cells from the polyclonal repertoire, we performed focused BCR repertoire analyses, demonstrating that accrual of affinity enhancing antibody mutations occurs readily for commensal-specific B cells following *de novo* colonization, and this does not result in polyreactive antibody responses, but elevated antibody binding affinities for this commensal antigen. Targeted isolation of antigen-specific B cells allowed a direct demonstration of gut B cell selection, and future studies to incorporate elegant photoactivation and fate-mapping approaches will enable us to extend these studies to the resolution of singular Peyer’s patches and even single germinal centers^31^.

Commensal-specific B cell tetramers will likely empower future investigation of the cellular heterogeneity in B cells responding to microbes in distinct experimental contexts, including heterologous immunity to commensal microbes during pathogen infection, malnutrition, autoimmunity, or within the tumor microenvironment. Key benefits of the B cell tetramer detection technology include the ability to precisely evaluate the cellular phenotype, isotype, kinetics and polyclonal repertoire of antigen-specific B cells recognizing an endogenous protein antigen expressed at physiological levels and kinetics. Additionally, B cell tetramers are compatible with fluorescent reporters and transcription factor staining, enabling ready integration into established flow cytometric panels and approaches. Importantly, because B cell tetramers are not subject to MHC restriction, they will enable comparisons of immunity to microbes across mouse haplotypes and across species, including humans and non-human primates.

In summary, here we developed an approach with which to identify immunogenic antigens from commensal microbes, and subsequently utilized identified antigens to track commensal-specific B cell responses *ex vivo*, with the view that this approach will facilitate future studies of humoral immunity to commensal microbes, throughout the lifespan, across generations, and with anatomical resolution.

### Limitations of the study

Utilization of phage-display library screening to identify immunogenic antigens is amenable for culturable microbes, or those that can be isolated with relative purity under gnotobiotic conditions. We have not systematically determined whether phage display libraries of sufficient size and complexity can be generated from diverse metagenomes to enable pan-commensal identification of immunogenic antigens. Generation of B cell tetramers is reliant upon stable expression of bacterial proteins in sufficient quantity to permit biotinylation and tetramerization, and as such, not all proteins may be amenable to being used as probes. Detection of antigen-specific B cells with tetramers is reliant on cell surface BCR/antigen interaction, currently negating the utility of these approaches to detect plasma cells due to low cell surface BCR expression. Ongoing efforts to generate commensal-specific BCR transgenic mice will likely circumnavigate these shortcomings.

## Materials and Methods

### Mice

Wild-type (WT) C57BL/6 Specific Pathogen Free (SPF) mice were purchased from Taconic and Jackson Laboratories. *Rag2*^−/−^/*Il2rg*^−/−^ (R2G2) mice were purchased from Envigo and colonized with SFB using feces from Taconic B6 mice. *Cd4*^cre^ x *Bcl6*^fl/fl^, *Icos*^−/−^ and *Tcrb*^−/−^*Tcrd*^−/−^ mice were generously provided by Marion Pepper (University of Washington), Daniel Campbell (Benaroya Research Institute), and Meghan Koch (Fred Hutchinson Cancer Center), respectively. All specified pathogen free (SPF) mice were maintained at Benaroya Research Institute, and experiments were approved by the Institutional Animal Care and Use Committee of Benaroya Research Institute. Germ-free mice and SFB-monocolonized mice were housed in the University of Washington Gnotobiotic Animal Core. All experimental procedures were approved by the Institutional Animal Care and Use Committee at the University of Washington. Mice used in experiments were between 6 and 12 weeks of age at time of sacrifice unless specified.

### Construction of phage display library

Genomic DNA was purified from the feces of SFB-monocolonized mice using DNeasy PowerSoil Pro Kit according to the manufacturer’s instructions. Purified DNA was fragmented by sonication using a Covaris M220 Focused-ultrasonicator. Fragmented DNA was analyzed on 1% agarose gels to ensure fragment sizes between 200 and 600 bp. Cohesive ends were blunted and repaired using Fast DNA End Repair Kit (Thermo Scientific). DNA was purified using a spin column (Zymo Research). The libraries were generated by blunt-end cloning of 1,200 ng fragmented DNA into 1,000 ng PmeI linearized and dephosphorylated pHORF3 library vector (a gift from Michael Hust, Technische Universität Braunschweig) (16 hr at 16 °C, T4 DNA Ligase, NEB). The ligation reaction was purified and transformed into TG1 bacteria (Lucigen) by electroporation (1.8 kV, MicroPulser, BioRad). After 1 hr of incubation at 37 °C and 600 rpm in 1 mL SOC medium, transformation rates were determined by plating dilutions on 2× YT agar (1.6% (w/v) tryptone, 1% (w/v) yeast extract, 0.05% (w/v) NaCl, 1.2% (w/v) agar) supplemented with 100 mM glucose and 100 μg/mL ampicillin (2× YT-GA). Cells were plated on 2× YT-GA agar plates (25 × 25 cm) and incubated at 37°C overnight. The cells were scraped using 20 mL of 2× TY medium, and stored at −80 °C in 20% (v/v) glycerol. Insert rates and mean insert sizes were determined by colony PCR and agarose gel electrophoresis of randomly analyzed colonies (n=24).

### Packaging of oligopeptide phage library

Four hundred mL of 2×TY-GA medium were inoculated with glycerol stock of library to an OD600 < 0.1 and incubated at 37 °C and 250 rpm until an OD600 of 0.5 was reached. In order to complement the missing coat proteins and ensure enrichment for ORFs, 25 mL of the culture were infected with a 20-fold excess i.e. 2.5 × 10^11^ colony forming units (cfu) of the M13K07ΔgIII helper phage “Hyperphage”(Progen) for 30 min at 37 °C. The infected cells were incubated for another 30 min at 37 °C and 250 rpm to express antibiotics resistance. The cells were pelleted (3,200× g, 10 min) and subsequently resuspended in 400 mL 2× YT medium supplemented with 100 μg/mL ampicillin and 50 μg/mL kanamycin (2× YT-AK). Oligopeptide phage particles were produced for 24 hr at 30 °C and 250 rpm. The supernatant containing phage particles were precipitated at 4 °C overnight after adding 1/5 volume precipitation buffer (20% (w/v) PEG 6,000, 2.5 M NaCl). The precipitated phage particles were pelleted for 1 h at 10,000× g and 4 °C (Sorvall Centrifuge) and resuspended in 10 mL of 1X PBS. A second precipitation step was performed for 1 h at 4 °C with 1/5 volume precipitation buffer. The phage particles were pelleted for 30 min at 20,000× g and 4 °C (Sorvall Centrifuge RC5B Plus, Rotor SS34) and resuspended in 1 mL 1X PBS. Remaining bacteria were pelleted for 2 min at 16100× g (Eppendorf Centrifuge 5415 D) and supernatants containing the oligopeptide phage libraries were collected and stored at 4 °C.

### Phage Library Biopanning

To reduce background signals, serum used for biopanning was precleared from antibodies reactive with phage particles. Therefore, two wells of a 96-well ELISA plate (Costar) were coated with 4 × 10^10^ cfu Hyperphage in 300 μL phosphate buffer saline (PBS). The protein binding capacity of the well surface was saturated with 300 μL blocking solution (PBS supplemented with 2% skim milk powder (BD) and 0.1% Tween 20) per well. The blocking solution was removed and the mouse sera from germ-free mice (GF) and SFB-monocolonized mice (SFB mono) (1:200 dilution in 300 μL blocking buffer) were incubated for 1 hr in each of the two wells (total 2 hr) with immobilized “Hyperphage” to preclear anti-phage serum antibodies.

Precleared serum from germ free and SFB monocolonized mice was transferred into 2 wells (150 μL each) previously coated with anti-mouse IgA and IgG antibodies and saturated with blocking buffer to capture the serum antibodies used for the panning procedure. Excess serum antibodies and other serum proteins were removed by washing 3 times with washing buffer (PBS supplemented with 0.05% Tween 20). For the selection of oligopeptides specific to germ-free serum antibodies, the phage library was incubated with immobilized GF serum antibodies for 1 hr at room temperature and then unbound phage were removed and placed onto wells containing serum from SFB monocolonized mice and further incubated for 2 hr at room temperature.

Unbound oligopeptide phage was removed by stringent washing (10 times with washing buffer). Bound oligopeptide phage particles were eluted with 200 μL elution buffer (10 μg/mL trypsin in PBS) per well for 30 min at 37 °C. The eluted phage titer was determined using 10-fold dilutions of 10 μL of the elution followed by *E. coli* TG1 infection and plating onto 2× YT agar supplemented with ampicillin. To amplify the eluted phage for input in the next panning round, the remaining 190 μL of elution were used to infect 1 mL of an *E. coli* TG1 culture (OD_600_ = 0.5) for 30 min at 37 °C. Cells were plated on 10 cm 2× YT-GA agar plates and incubated at 37 °C overnight to allow amplification of eluted phage for the next panning round. The cells were scraped using 5 mL of 2× YT-GA. To produce phage, 30 mL of 2× YT-GA were inoculated (OD_600_ < 0.1) with the scraped cells and grown up to OD_600_ = 0.5. Phage particles were produced following the same procedure as described above for packaging of oligopeptide phage libraries. Amplified phage was precipitated only once (1 hr on ice) after adding 1/5 volume precipitation buffer. After pelleting (1 hr at 4 °C, 3,220× g), the phage was resuspended in 1000 μL of 1X PBS. Remaining bacteria were pelleted for 10 min at 15000× g, the amplified oligopeptide phage containing supernatant was collected and stored at 4 °C until used as input phage for the next panning round. After the fourth panning round, 10-fold dilutions of eluted phage were used to infect 50 μL *E. coli TG1* (OD_600_ = 0.5), the cells were plated on 2× YT-GA agar plates and incubated at 37 °C overnight to obtain single colonies that allow screening of monoclonal oligopeptide phage. To screen the enriched clones after the third round of each panning for immunogenic oligopeptides, 96 randomly selected clones were analyzed by monoclonal phage ELISA.

### Monoclonal Phage ELISA

To screen clones obtained from biopanning monoclonal phages were produced. A 96-well microtiter plate was supplemented with 180 μL 2× YT-GA medium, inoculated with single colonies after the fourth panning round and incubated (37 °C, 300 rpm) overnight. For phage production, 175 μl 2× YT-GA per well were inoculated with 10 μL of the overnight culture and incubated for 2 hr at 37 °C and 300 rpm to reach logarithmic growth. Cells were infected with 5 × 10^9^ cfu Hyperphage (M13K07ΔgIII) for 30 min at 37 °C. To ensure antibiotic resistance, cells were incubated for 30 min at 37 °C and 300 rpm. To change the medium for phage production, the cells were pelleted for 10 min at 3,220× g and resuspended in 180 μL 2× YT-AK followed by phage production at 30 °C and 300 rpm overnight. Cells were pelleted and supernatant containing phages used for monoclonal phage ELISA.

To capture phage, rabbit anti-M13 (pVIII specific, 1:5,000 in PBS, PA1-26758 Thermo Scientific) was immobilized on an ELISA plate (Costar) at 4 °C overnight. Wells were saturated with blocking buffer for 1 hr at room temperature. Plates were washed 3 times with washing buffer before addition of monoclonal phage (50 μL in 100 μL blocking buffer). Phage particles were captured for 2 hr at room temperature. To reduce background phage binding, mouse serum samples (1:500) were precleared in blocking buffer containing 10^10^ cfu/mL Hyperphage for 2 hr at room temperature. Remaining monoclonal phage were removed by 3 washing steps with washing buffer and the precleared sera were transferred to the wells containing immobilized oligopeptide phage particles and incubated for 2 hr at room temperature. Serum was removed by 3 washing steps and bound serum antibodies were detected using goat anti-mouse IgA and IgG antibodies conjugated with horseradish peroxidase (HRP) for 1 hr at room temperature. Excess detection antibody was removed by 3 washing steps and the ELISA developed with TMB substrate solution. Signals were detected using an ELISA reader at 450 nm using SpectraMax M2.

### Enzyme-linked immunosorbent assay (ELISA)

High binding ELISA plates (Corning) were coated overnight at 4 °C with 2.5 ug/ml of SFB3340 protein. Plates were blocked with 2% skim milk in PBS prior to sample incubation. For serum samples, plates were incubated with serially diluted serum. For cloned mAbs, plates were incubated with serially diluted mAbs starting at 10 ng/ml. Each sample was plated in duplicate. For serum samples, bound antibodies were detected using peroxidase conjugated goat antibodies to IgA (SouthernBiotech), IgG_1_ (SouthernBiotech), IgG_2b_ (SouthernBiotech), IgG_2c_ (Jackson Lab), IgM (Jackson Immunoresearch) at 1:5000 dilution in PBS. For mAbs, bound antibodies were detected with mouse anti-human IgG-HRP. Absorbance was measured at 450 nm using SpectraMax M2.

### ELISpot

Multiscreen 96-well ELISPOT plates (Millipore) were coated overnight at 4 °C with 2.5 ug/ml of SFB3340 protein. Plates were blocked with 10% FBS in complete RPMI (GIBCO). Cells from various tissues were serially diluted in complete RPMI and plated onto coated ELISPOT plates and incubated at 37°C overnight. Cells were washed off and following washes with PBS, secondary peroxidase conjugated goat antibodies to IgG_1_, IgG_2b_ and IgA (Southern Biotech) were used at 1:1000 in PBS to detect antibody-secreting cells. Plates were developed with AEC developing reagent (Vector Laboratories) according to manufacturer’s instructions. Plates were read on an ImmunoSpot C.T.L. Analyzer and quantitated using ImmunoSpot.

### Tetramer generation

Recombinant SFB3340 protein was procured from GenScript, biotinylated at 1:1 ratio using EZ-Link™ Sulfo-NHS-LC-Biotin (ThermoFisher) and tetramerized with streptavidin-PE (Prozyme) as previously described^42^. Decoy reagent to gate out non-SFB3340^+^ cells was made by conjugating SA-PE to DyLight™ 594 NHS Ester (ThermoFisher), washing and removing any unbound DyLight™ 594, and incubating with an excess of an irrelevant biotinylated decoy peptide bearing the affinity purification tags present in SFB3340 protein used for tetramer generation, specifically, a 31 mer peptide (HHHHHHENLYFQGGSGGLNDIFEAQKIEWHE) containing His tag, TEV cleavage sites and AviTag sequence was used to make the decoy reagent.

### De novo colonization with SFB containing feces

For SFB colonization, fresh fecal pellets collected from *Rag2*^−/−^*Il2rg*^−/−^ mice were resuspended in PBS and clarified by centrifuging the bacterial pellet at 300g, prior to oral gavage.

### Quantification of fecal DNA

Fecal DNA was prepared using the Quick-DNA Fecal/Soil Microbe DNA Miniprep Kit (Zymo Research) according to the supplier’s protocol. For generating a standard curve of qPCR, SFB 16S rRNA gene fragment was cloned into the pCR™2.1-TOPO™ vector (ThermoFisher) using TA cloning. A serial 10-fold dilution of 10^2^, 10^3^, 10^4^, 10^5^, 10^6^, 10^7^, 10^8^ and 10^9^ copies of the recombinant plasmid DNA containing the SFB 16S rRNA gene fragment was used. Two pairs of SFB 16S rRNA gene-based qPCR primers, SFB736F (5′-GACGCTGAGGCATGAGAGCAT-3′)/SFB844R (5′-GACGGCACGGATTGTTATTCA-3′) were used in a 7500 Fast Real-Time PCR System (Life Technologies) to quantify SFB level in the feces. Briefly, a 10-μl mixture containing 50ng of template, 1μl of the primer mixture containing 100 nM of each of the forward and reverse primers, 5μl of 2 x SYGR green Fast master mix (Life Technologies) were used to set up the reaction.

### Flow cytometry

Single-cell suspensions were prepared and resuspended in 50μl in PBS containing Fc block (2.4G2) and first incubated with decoy tetramer at a concentration of 10 nM at room temperature for 10 min. SFB3340-PE tetramer was added at a concentration of 10 nM and incubated on ice for 30 min along with surface antibodies followed by intracellular antibody staining when needed. All cells were run on a BD Symphony A5 cytometer and analyzed using FlowJo software.

### Isolation of leukocytes from tissues

Cells from Peyer’s patches, mesenteric lymph nodes, spleen and bone marrow were collected in complete media (RPMI, 3% heat inactivated FCS, 10 mM HEPES, 1% Pen/Strep). The tissue was mashed through a 70μm cell strainer to generate single cell suspensions. For lamina propria, the small intestine was dissected and opened longitudinally and cut into 4-cm pieces, followed by incubation in RPMI, 3% heat inactivated FCS, 5 mM EDTA (Sigma-Aldrich), 0.145 mg/ml DTT (Sigma-Aldrich) at 37°C for 20 minutes on a stirring platform. To separate the intestinal epithelial lymphocyte (IEL) and lamina propria (LP) layer, intestinal contents were transferred through a sterile fine-meshed kitchen strainer into a 500 ml beaker. Strainer was tapped on beaker several times after straining and pieces of small intestine were transferred to a 50 ml conical tube containing 10 ml of serum free media with 2mM EDTA per intestine. After shaking the tube vigorously for 30 seconds, the contents of the tube were strained through the strainer into the beaker. The process was repeated three times. The tissue was minced finely in the digest containing RPMI with 0.1 mg/ml liberase CI (Roche) and 1mg/ml DNase I (Sigma-Aldrich), with continuous stirring at 37°C for 25 min. Digested tissue was passed through 100-um cell strainers. Lymphocytes were further enriched by centrifugation at room temperature at 695 g for 8 min in 37.5% Percoll (GE Healthcare), prior to washing and resuspension for downstream analysis.

### Single-cell RNA-sequencing

Peyer’s patch leukocytes from 10 WT B6 mice de novo colonized for 21 days with SFB were separately stained with oligo-hashtagged antibodies (TotalSeqC 1 – 10, Biolegend), prior to pooling, staining, and FACS isolation of SFB3340^+^ B cells. A single cell suspension was prepared from pooled sorted cells and loaded onto one channel on the 10x Chromium Controller (10x Genomics) according to the manufacturer’s protocol, with a target capture of 10,000 cells per channel. Sequencing libraries were generated using the NextGEM Single Cell 5’ Kit v2. Gene expression, BCR, and feature barcoding libraries were pooled and treated with Illumina Free Adapter Blocking Reagent (Illumina). Sequencing of pooled libraries was carried out on a NextSeq 2000 sequencer (Illumina), using a NextSeq P2 flowcell (Illumina), with target depths of 40000, 5000, and 5000 raw reads per cell for gene expression, BCR, and feature barcode libraries, respectively. Basecalls were converted to FASTQs and processed to gene counts, hashtag counts, and assembled BCR sequences using Cell Ranger v. 6.1.1, with the GRCm38 genome annotation and V(D)J reference, using default parameters and cell calling with an expected cell number of 10000.

### Single-cell RNA-sequencing quality control and demultiplexing

Filtering, normalization, projection, and clustering of 10x data was conducted using Seurat Toolkit v.4.3^52^. Cell Ranger processing yielded 2984 10x barcodes called as putative cells. To limit analysis to high-quality single cells, quality filters were applied to the gene count data using the following empirical thresholds: total genes detected ≥ 250 and ≤ 6000, UMI counts ≥ 1400 and ≤ 35000, mitochondrial UMIs ≤ 20%, ribosomal protein UMIs ≤ 50%, and hemoglobin UMIs ≤ 10%. To limit analysis to B cells, we excluded barcodes expressing genes characteristic of other cell types (raw UMI counts: Cd3d ≥ 3, Cd3e ≥ 3, Vil1 ≥ 1, Reg3g ≥ 2), and retained only barcodes expressing characteristic B cell genes (log-normalized UMI counts: Cd74 ≥ 2, H2-Ab1 ≥ 0.5, Cd79a ≥ 0.5, Ly6e ≥ 0.5, Cd52 ≥ 0.25). We used empirical thresholds on the log-counts-per-million hashtag UMIs to call each barcode as positive or negative for each hashtag, and kept cells that were positive for a single hashtag. After quality filtering and demultiplexing, we retained 1762 high-quality single B cells for downstream analysis.

Gene expression data were log-normalized, projected onto a UMAP embedding, and clustered using shared nearest neighbor modularity optimization as implemented in Seurat. Cluster identity was determined using marker gene expression, and confirmed using Ig isotypes based on heavy chain alignments in the assembled BCR sequences.

### Analysis of BCR sequence data

We analyzed BCR sequence data using the Immcantation toolkit (https://immcantation.readthedocs.io/en/stable/). Starting from the assembled BCR contig FASTA files output by cellranger, Ig regions and gene usage for each sequence were determined using Change-O^53^. BCR clones were identified independently for heavy and light chains using Change-O, using optimal clone distances estimated with Shazam^53^. Heavy chain clones were filtered to include only those with consistent light chain sequences, and germline sequences were inferred at the nucleotide level. BCR lineages were estimated from the full amino acid sequences using the parsimony ratchet^54^, as implemented in the R package phangorn^55^. We used custom R scripts to determine mutations from the germline sequence at the nucleotide level.

### Recombinant antibody production

Recombinant antibodies derived from recurrent public clonotypes were generated by WuXi Biologics in a human IgG_1_ backbone.

### Site directed mutagenesis (SDM)

SDM was carried out using germline sequences from two public (VH1-76(RC) and VH1-53(RC)) clonotypes as DNA template. Mutagenic primers were designed using PrimerX. Briefly, 50 ng of template DNA, 0.5 µM forward and reverse primer, Phusion High–Fidelity DNA Polymerase (Thermo Scientific) were used, and annealing temperature was set to 58°C. After PCR, the reaction was treated with 1µl of DpnI (New England Biolabs) to degrade parental DNA. The digested DNA sample was purified using a spin column (Zymo Research) and transformed into chemically competent Stbl3 cells. The colonies obtained were verified using Sanger sequencing.

### Recombinant antibody production for SDM

Expi293 cells were cultured in Expi293 Expression Medium following manufacturer’s instructions. (Thermo Scientific). Cells were transfected with 625 ng of IgH and corresponding IgL chain vector DNA in 24-well deep-well plates using the Expi293 Transfection Kit. Cells were grown at 37°C, 8% CO2, and 250 rpm for 6 days. On day 6 post transfection, the culture supernatant was harvested and loaded on a protein G column. The column was washed with PBS, and the IgG protein was eluted with a low pH buffer followed by buffer exchange in 1X PBS.

### Biolayer Interferometry

All assays were performed on the Octet Red Instrument (ForteBio) at 30C with shaking at 1,000 RPM. Purified antibodies were captured using Anti-Human IgG Fc capture (AHC) biosensors by immersing sensors into KB buffer (1X PBS, 0.01% BSA, 0.02% Tween 20, and 0.005% NaN3) containing individual antibodies at a standard concentration of 10ug/mL for 300s. After loading, baseline signals were recorded for 60s in KB. The sensors were then immersed in wells containing SFB3340 at a standard concentration of 250nM in KB for 300s (association phase). Following this, sensors were immersed in KB for an additional 300s (dissociation phase). Rmax values were recorded as the max nm shift at the end of the association phase. All binding experiments were run twice for each antibody, and mean Rmax values were reported.

### Immunofluorescence staining

Ileal tissue R2G2 mice were flushed with cold PBS, splayed open and rolled into a Swiss roll, then fixed in 4% paraformaldehyde (PFA) for 4hr and placed in 30% sucrose overnight at 4°C. Next day rolls were shaken gently in OCT for 2hr and then embedded in OCT.16 μm sections were cut on a cryotome, dried and then frozen at −80°C. Tissues were blocked and permeabilized using 5% Rat IgG, 5% BSA, 0.3% Triton-X in PBS for 1 hr. Tissues were stained with Alexa Fluor® 488 anti-mouse EpCAM (1:200 dilution in 5% BSA in PBS) antibody and recombinant mAb V_H_1-76RC N1 overnight at 4°C. Anti-human IgG1-R718 (1:100 dilution in 5% BSA in PBS) was used as a secondary antibody. Images were acquired using an SP5 confocal microscope (Leica).

## Acknowledgements

We thank members of the Harrison Lab for discussion and critical reading of the manuscript. We thank Marion Pepper for *Cd4*^Cre^ x *Bcl6*^fl/fl^ mice, Dan Campbell for *Icos*^−/−^ mice and Meghan Koch for *Tcrbd*^−/−^ mice. We thank Justin Taylor for technical guidance on B cell tetramer generation and usage. We also thank Benaroya Research Institute Cell and Tissue Analysis and Genomics Cores, particularly Caroline Stefani, Adam Wojno, Vivian Gersuk, Kimm O’Brien and Quynh-Anh Nguyen, as well as staff of the Benaroya Research Institute vivarium. We thank Jisun Paik for overseeing gnotobiotic experiments. We thank Michael Hust for provision of the pHORF3 vector, and Dan Littman for sharing SFB monocolonized feces. We thank the M.J. Murdock Charitable Trust for support of flow cytometry, histology, imaging and genomics resources at Benaroya Research Institute.

## Author Contributions

S.V. carried out most of the experiments and analyzed the data. M.D. performed analysis of single-cell RNA-sequencing and BCR repertoires. T.M.O., S.K., and J.L., aided with *in vivo* experiments and analyses. S.S., and A.T.M., performed antibody binding and affinity experiments and analyses. O.J.H conceived the project, analyzed data and wrote the manuscript with input from all co-authors.

## Lead contact and materials availability

All correspondence and material requests should be addressed to O.J.H.

**Supplementary Figure 1:** (A) Schematics showing 4 immunogenic proteins and targeted epitopes identified by SFB-specific phage library screening. SFB3340-specific ELISA of serum antibody reactivity by (B) Tac and Jax B6 mice, (C) WT and *Tcrbd*^−/−^ mice, (D) WT and Icos^−/−^ mice and (E) *Bcl6*^ΔCD4^ and *Bcl6*^fl/fl^ littermates. (F) Representative images and quantification of SFB3340-specific ELISpot IgG_1_ and IgG_2b_ responses from leukocytes isolated from indicated tissues and stimulated with recombinant SFB3340. Data represent one of two independent experiments with three to six mice per group.

**Supplementary Figure 2:** (A) Schematic of protein domains in SFB3340 tetramer and decoy reagents. (B) Representative gate of B220 and CD138 expression on lineage negative lymphocytes prior to tetramer and decoy gating.

**Supplementary Figure 4:** (A) Quantification of SFB abundance in stool pellets from mice *de novo* colonized with SFB-containing fecal contents from *Rag2*^−/−^*Il2rg*^−/−^ donors, determined by qPCR at indicated days post colonization.

**Supplementary Figure 5:** (A) UMAP visualization of B cell clusters obtained by demultiplexing single-cell RNA sequencing data from individual mice based on hashtag antibodies. (B) ELISA-based binding of monoclonal antibody obtained from two clonotypes V_H_1-53RC-V_k_5-43J_k_2 and V_H_1-76RC-V_k_5-48J_k_2 against bovine serum albumin, double-stranded DNA, lipopolysaccharide, and ovalbumin.

