## Supplementary figures and images for "Antigen-level resolution of commensal-specific B cell responses enabled by phage-display screening and B cell tetramers"

### Supplementary Figures

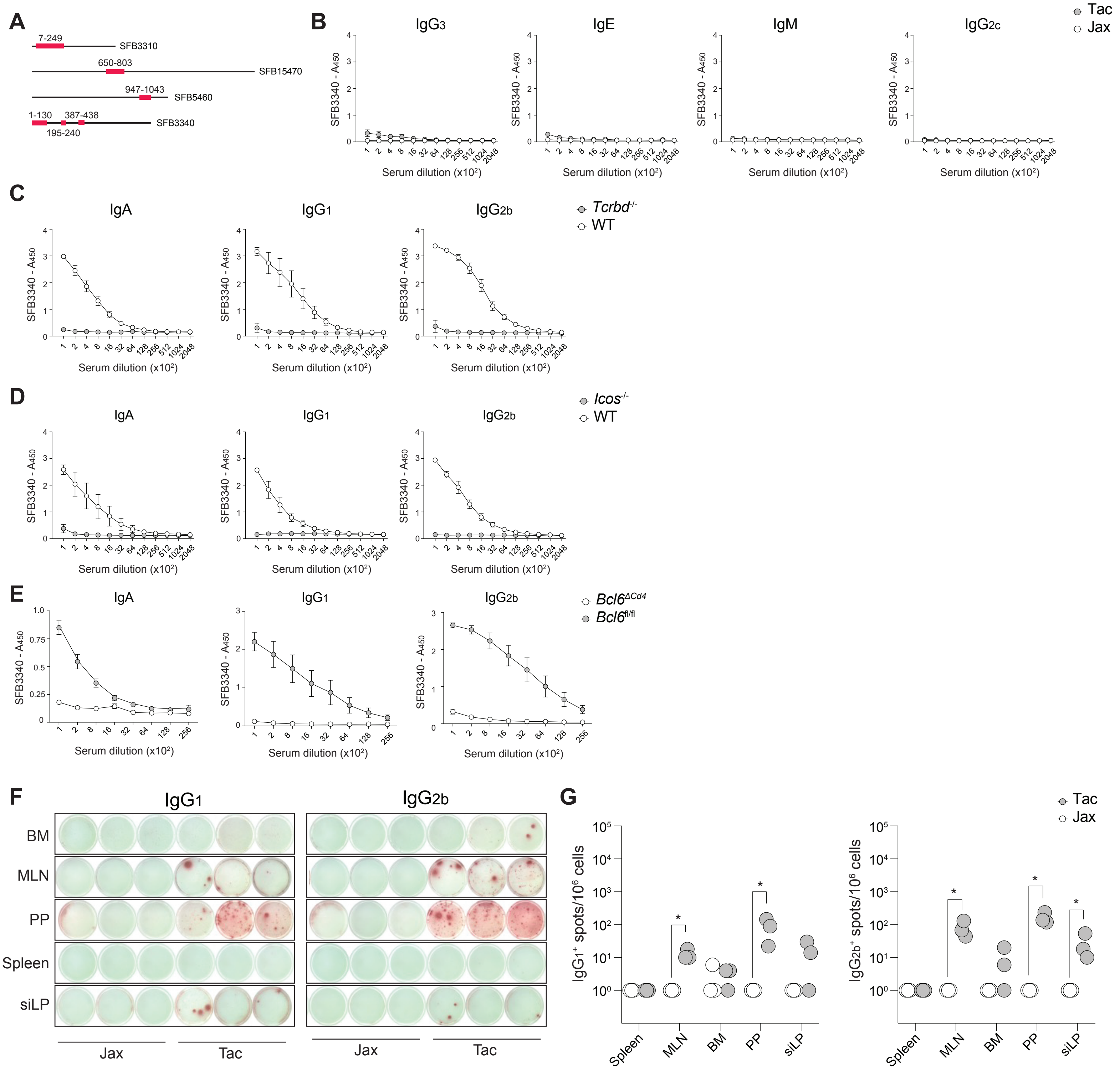


Verma *et al* - Figure S1

**A**

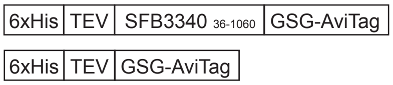


**B**

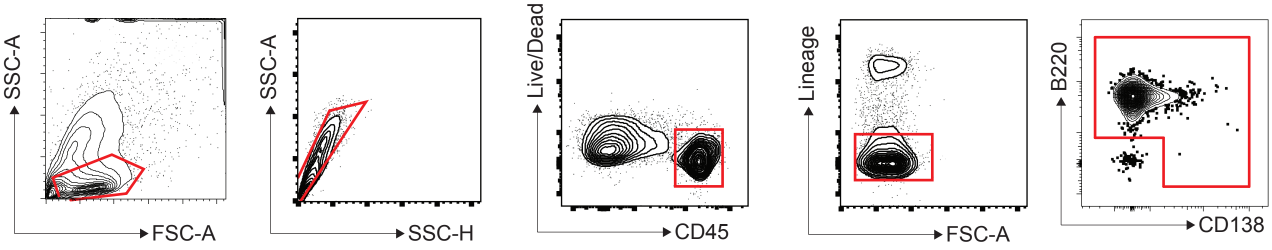


Verma *et al* - Figure S2

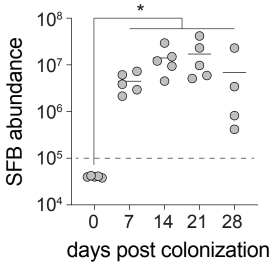

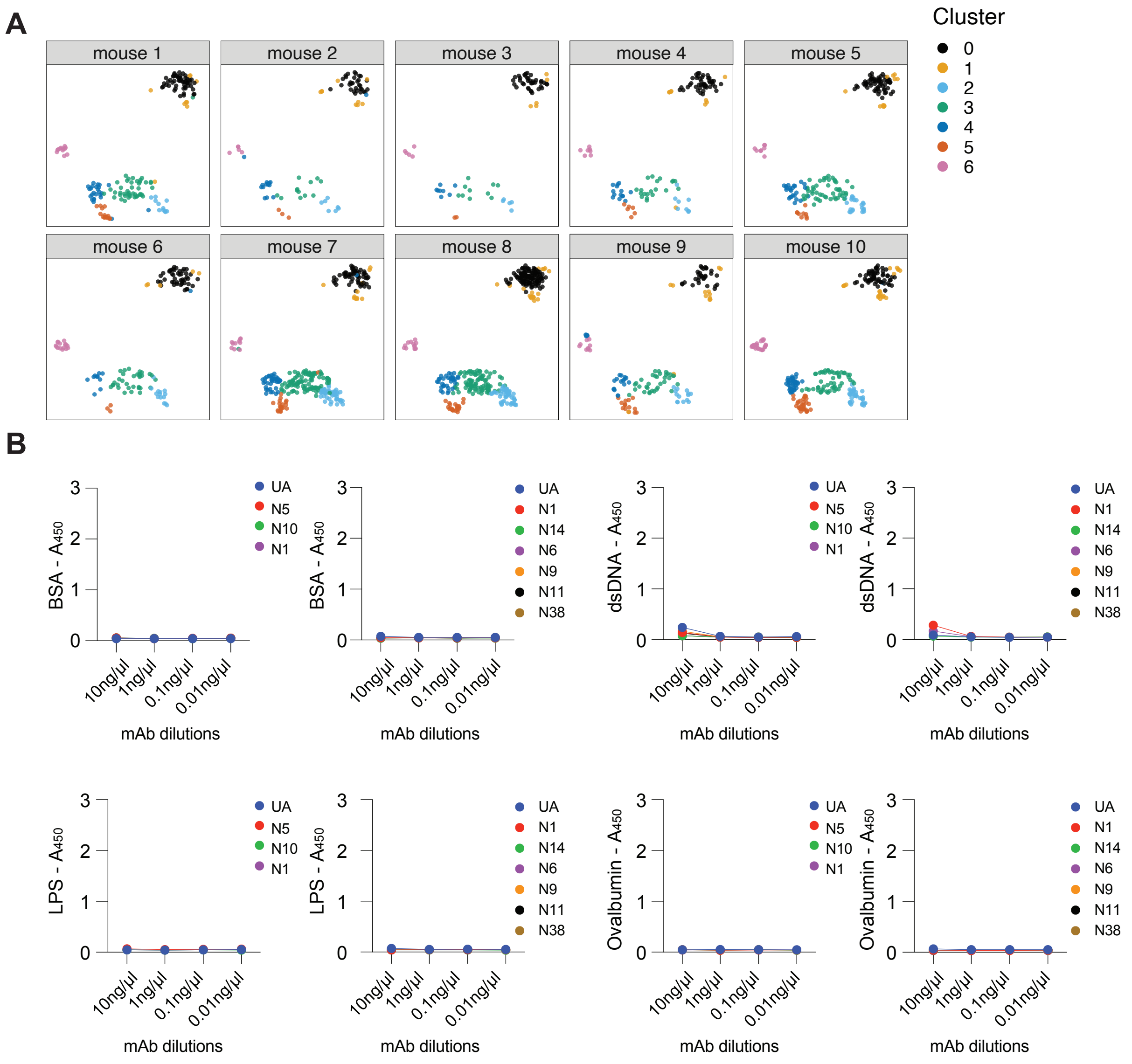


Verma *et al* - Figure S5
